# Bone Morphogenetic Protein signaling pathway – ethanol interactions disrupt palate formation independent of *gata3*

**DOI:** 10.1101/2024.11.15.623833

**Authors:** C. Ben Lovely

## Abstract

Fetal Alcohol Spectrum Disorders (FASD) describes a wide array of neurological defects and craniofacial malformations, associated with ethanol teratogenicity. While there is growing evidence for a genetic component to FASD, little is known of the genes underlying these ethanol-induced defects. Along with timing and dosage, genetic predispositions may help explain the variability within FASD. From a screen for gene-ethanol interactions, we found that mutants for Bmp signaling components are ethanol-sensitive leading to defects in the zebrafish palate. Loss of Bmp signaling results in reductions in *gata3* expression in the maxillary domain of the neural crest in the 1st pharyngeal arch, leading to palate defects while upregulation of human *GATA3* rescues these defects. Here, we show that ethanol-treated Bmp mutants exhibit misshaped and/or broken trabeculae. Surprisingly, up regulation of *GATA3* does not rescue ethanol-induced palate defects and *gata3* expression was not altered in ethanol-treated Bmp mutants or dorsomorphin-treated larvae. Timing of ethanol sensitivity shows that Bmp mutants are ethanol sensitive from 10-18 hours post-fertilization (hpf), prior to Bmp’s regulation of *gata3* in palate formation. This is consistent with our previous work with dorsomorphin-dependent knock down of Bmp signaling from 10-18 hpf disrupting endoderm formation and subsequent jaw development. Overall, this suggests that ethanol disrupts Bmp-dependent palate development independent of and earlier than the role of *gata3* in palate formation by disrupting epithelial development. Ultimately, these data demonstrate that zebrafish is a useful model to identify and characterize gene-ethanol interactions and this work will directly inform our understanding of FASD.

**Highlights:**

- Bmp pathway mutants are ethanol sensitive resulting in palate defects.
- Ethanol disrupts Bmp-dependent palate development independent of *gata3*.
- Timing of ethanol sensitivity suggests ethanol disrupts Bmp-dependent epithelial morphogenesis.

## Introduction

Fetal Alcohol Spectrum Disorders (FASD) describe the most common, preventable cause of congenital anomalies. Though worldwide incidence rates of FASD are .77%, the incidence rates in the US range from 2-5%, with incidence rates in other parts of the world topping nearly 30% due to extreme rates of binge drinking (May et al., 2018; Popova et al., 2019). However, these are likely underestimates as 1) up to 10% of women worldwide consume alcohol during pregnancy, 2) nearly 50% of pregnancies are unplanned, and 3) many pediatricians fail to recognize and diagnose FASD (Finer and Zolna, 2016; Popova et al., 2018, 2017; Rojmahamongkol et al., 2015). FASD is highly variable presenting with multiple phenotypes, including structural malformations to the brain and face (Lovely, 2020). At the most severe end of the FASD spectrum is Fetal Alcohol Syndrome (FAS), which presents with and is diagnosed by craniofacial defects including deficits in midfacial depth, philtrum morphology, and formation of Oral-facial cleft (OFCs) (Beaty et al., 2011; Bemquerer et al., 2022; Blanck-Lubarsch et al., 2020; Suttie et al., 2017, 2013). However, it is becoming increasingly clear that children exposed prenatally to ethanol can have craniofacial defects in the absence of a diagnosis of FAS (Muggli et al., 2017; Suttie et al., 2013).

A significant barrier to our understanding of FASD is a lack of insight in the genetic predisposition to ethanol-induced developmental defects. Several lines of evidence in multiple vertebrates, including human, suggest a genetic component to the prevalence and variability of FASD phenotypes. The most direct evidence of genetic risk factors in humans come from twin studies that demonstrate 100% concordance in FASD phenotypic outcomes in monozygotic twins over dizygotic twins (66%), full siblings (41%) and half siblings (22%) (Hemingway et al., 2018). However, the complex interplay between genetic background and timing and dosage makes studying FASD in humans challenging. Therefore, genetically tractable model organisms have been essential in improving our understanding of the genetic risk factors in FASD (Eberhart and Parnell, 2016; Lovely, 2020). To date, several ethanol-sensitizing alleles in in zebrafish, mouse and humans have been linked with increased cell death, holoprosencephaly, oral clefting, disruption to axonal projections, and broad neural and eye defects (Boyles et al., 2010; Hong and Krauss, 2012; McCarthy et al., 2013; Swartz et al., 2014; Zhang et al., 2013). However, despite these growing insights into the genetic components contributing to risk for FASD, we lack insight into ethanol-sensitive gene function and associated cellular mechanisms during development that underlie FASD (Lovely, 2020).

Here, we use zebrafish to improve our understanding of these ethanol-sensitive mechanisms behind craniofacial phenotypes in FASD. The zebrafish is perfectly suited for studying ethanol-sensitizing alleles and pathways with its genetic tractability, high fecundity, external fertilization, embryo transparency, and rapid development and work in zebrafish does translate to studies of candidate genes in human populations (Klem et al., 2023 (in revision); McCarthy et al., 2013). With this high degree of genetic and molecular conservation, zebrafish studies have played a pivotal role revealing the mechanisms underlying facial development. We have previously used several genetic-based approaches to identify ethanol-sensitive mutations (Klem et al., 2023 (in revision); McCarthy et al., 2013; Swartz et al., 2020, 2014) and these approaches have proven successful in predicting human gene-ethanol interactions (Klem et al., 2023 (in revision); McCarthy et al., 2013). However, despite the increasing number of identified ethanol-sensitive loci (Beaty et al., 2011; Boyles et al., 2010; Hong and Krauss, 2012; Kietzman et al., 2014; Langevin et al., 2011; McCarthy et al., 2013; Swartz et al., 2020, 2014; Zhang et al., 2013, 2011), our mechanistic understanding of the impact of these gene-ethanol interactions on the complex processes underlying craniofacial development is lacking.

Formation of the facial skeleton is a complex process involving cells from all three germ layers in particular ectoderm derived neural crest cells (NCCs). Highly conserved amongst all vertebrates the NCCs migrate to homologous regions of the forming head to form the majority of the craniofacial skeleton (Cordero et al., 2011; Santagati and Rijli, 2003). Through temporally and genetically conserved molecular mechanisms, the highly homologous bones constituting the facial skeleton including the palate take shape (Machado and Eames, 2017). The NCCs that form the palate migrate to populate the maxillary domain of the 1st pharyngeal arch, which lies just ventral to the developing eye but dorsal to the oral ectoderm (Cordero et al., 2011; Santagati and Rijli, 2003). The NCCs in the 1st pharyngeal arch undergo reciprocal signaling interactions with the adjacent facial epithelia, in particular the anterior endoderm and oral ectoderm (Choe and Crump, 2015; Mork and Crump, 2015). These signaling interactions are critical to the morphogenesis of the NCCs and subsequent palate formation (Swartz et al., 2011). The palate in zebrafish is comprised of an anterior, midline ethmoid plate which fuses to two bilateral cartilage rods called trabeculae (Eberhart et al., 2006; Swartz et al., 2021). These trabeculae then fuse with the posterior neurocranium at the polar cartilages (McCarthy et al., 2016). Overall the ethmoid plate and trabeculae are analogous to the mammalian palate with a high degree of genetic conservation in palate formation between zebrafish and mammals (Swartz et al., 2011).

Multiple studies have demonstrated that loss-of-function in Bmp pathway members results in cleft palate in mice (Dudas, *et al*. 2004; Liu, *et al*. 2005; Ko, *et al*. 2007). Bmp signaling is a complex pathway essential for craniofacial development and work in zebrafish has shown that Bmp signals directly to the NCCs for palate morphogenesis regulating multiple targets including *gata3* (Alexander et al., 2011; Sheehan-Rooney et al., 2013; Swartz et al., 2021, 2011). Mutations in human *GATA3* result in the highly variable hypoparathyroidism, sensorineural deafness and renal dysplasia (HDR) syndrome, which is also associated with palatal defects (Bilous et al., 1992; Van Esch et al., 2000). We have previously demonstrated that zebrafish *gata3* mutants lose a specific region of the palatal skeleton, the trabeculae, phenocopying loss of Bmp signaling (Alexander et al., 2011; Swartz et al., 2021) and that knock down of Bmp signaling with the small chemical inhibitor, dorsomorphin (DM), from 10-18 hpf disrupts endoderm formation and subsequent jaw development (Lovely et al., 2016). Recently, we have shown that mutation in the Bmp ligands, *bmp2b* and *bmp4* and the receptor *bmpr1bb* (defined as Bmp mutants going forward) sensitize larvae to ethanol-induced jaw defects by disrupting anterior endoderm morphology (Klem et al., 2023 (in revision)) suggesting that these Bmp mutants may also exhibit ethanol-induced palate defects. Here, I tested the ethanol sensitivity of these Bmp mutants during palate development. I show that all three mutants exhibit a range of defects to the palate, from misshapen trabeculae to breaks at the joint between the trabeculae and polar cartilages. I show that ethanol-induced disruptions to palate formation are independent of *gata3* with upregulation of human *GATA3* failing to rescue ethanol-induced palate defects in the Bmp mutants. Expression of *gata3,* which we have previously shown to be Bmp dependent, is unaffected in ethanol-treated Bmp mutants. I go on to show that our Bmp mutants are ethanol sensitive from 10-18 hpf, consistent with DM-dependent Bmp knockdown, suggesting that the Bmp mutants are ethanol sensitive before Bmp’s role in *gata3* dependent palate formation. Collectively, our data demonstrates that despite our growing insight into the ethanol-sensitive genetic loci in FASD, we still lack the mechanistic insights into the ethanol-sensitive gene function and subsequent cell mechanisms underlying FASD.

## Materials and Methods

### Zebrafish (*Danio rerio*) care and use

All zebrafish were raised and cared for using established IACUC protocols approved by the University of Louisville (Westerfield, 2007). Adult fish were maintained at 28.5°C with a 14 / 10-hour light / dark cycle. The *bmp2b^tc300a^* (Mullins et al., 1996), *bmp4^st72^* (Stickney et al., 2007), *bmpr1bb^sw40^* (Neumann et al., 2011) and *hsp70:GATA3-EGFP^au34^* (Swartz et al., 2021) zebrafish lines were previously described. Sex as a biological variable is not applicable at our studied development stages as sex is first detectable in zebrafish around 20-25 days post-fertilization (Aharon and Marlow, 2022), after all of our analyses.

### Zebrafish staging and ethanol or dorsomorphin treatment

Eggs from random heterozygous crosses were collected and embryos were morphologically staged (Westerfield, 2007), sorted into sample groups of 100 and reared at 28.5°C to desired developmental time points. All groups were incubated in embryo media (EM). Embryo media is a buffered solution at a pH of 7.2 that was modified from Westerfield, 2007. EM consists of a stock solution that is comprised of 17.5g NaCl, 0.75g KCl, 2.9g CaCl_2_, 0.41g K_2_HPO_4_, 0.142g NA_2_HPO_4_, 4.9g MgSO_4_ · 7H_2_O dissolved in 1 L of water, filter sterilized and stored at 4°C. To generate EM working solution, combine the 1 L of EM stock solution and 1.2g NaHCO_3_ and fill to 20 L with water and store at room temperature. At 10 hpf, EM was changed to either fresh EM or EM containing 1% ethanol (v/v), 10 µM dorsomorphin (DM) or an equivalent volume of DMSO. The end of the exposure window was based on the experimental design but EM containing ethanol, DMSO or DM was washed out with three fresh changes of EM.

### Cartilage and Bone staining

Zebrafish larva fixed at 5 dpf were stained with alcian blue for cartilage and alizarin red for bone (Walker and Kimmel, 2007). The palate was dissected free of surrounding tissues with fine stainless steel minutien pins (FST) and flat mounted for imaging (Kimmel et al., 1998). Brightfield images of the palate were taken on an Olympus BX53 compound microscope.

### Hybridization chain reaction (HCR) *in situ* hybridization

Embryos were collected at 36 hpf, dechorionated and fixed in 4% paraformaldehyde/PBS at 4°C. HCR protocol was previously described (Ibarra-García-Padilla et al., 2021). HCR amplifiers and buffers were acquired from Molecular Instruments (Molecular Instruments). The HCR probe against *gata3* was designed as previously described (Kuehn et al., 2022). Lateral confocal images of *gata3* expression at 36 hpf were taken using an Olympus FV1000 microscope and processed in FIJI (Schindelin et al., 2012).

### Statistical analyses

Palate defects in the single Bmp mutants were analyzed with a Fisher’s Exact Test. Palate defects in *hsp:GATA3-GFP* rescue experiments were analyzed with a two-way ANOVA (type III) with a Tukey’s Multiple Comparisons Test in Graphpad Prism 9.5.1 (Graphpad Software Inc., La Jolla, CA).

## Results

### Mutations in multiple components of the Bmp pathway sensitize embryos to ethanol-induced palatal defects

We previously screened several components of the Bmp signaling pathway (ligands *bmp2b* and *bmp4* and the receptor *bmpr1bb*) for ethanol sensitivity and showed ethanol-induced disruption to jaw development (Klem et al., 2023 (in revision)). In addition to jaw malformations, we also observed defects in palatal skeleton. In this screen, we exposed these Bmp signaling mutants to a sub-phenotypic dose of 1% ethanol (v/v) from 6 hpf to 5 days post fertilization (dpf) as this is highest dose that does not cause craniofacial defects in wild type embryos (Bilotta et al., 2004; Everson et al., 2022; McCarthy et al., 2013; Swartz et al., 2014; Zhang et al., 2014). This results in an average ethanol tissue concentration of 50 mM (approximately 30% of the media) (Flentke et al., 2014; Lovely et al., 2014; Reimers et al., 2004; Zhang et al., 2013) and is roughly equivalent to a human Blood Alcohol Concentration of 0.23. Though a binge dose, this is physiologically relevant to FASD with humans readily surpassing this amount (Canfield et al., 2019; Ethen et al., 2009; Jones, 2008; Maier, 2001; Whaley et al., 2019).

Using this dose, I found that mutations in our Bmp components sensitize larvae to a range of palate malformations. Indistinguishable from their wild type siblings, untreated heterozygous *bmp2b* or heterozygous and homozygous mutant *bmp4* and *bmpr1bb* larvae undergo normal craniofacial development (*bmp2b* mutant embryos do not develop past 16 hpf (Nguyen et al., 1998)) (Fig. 1A-C, J). However, when exposed to 1% ethanol (v/v), these Bmp mutants developed a range of palate malformations from misshapen trabeculae (the bilateral cartilage rods linking the posterior neurocranium with the ethmoid plate) to breaks in the trabeculae at or near the polar cartilages (posterior neurocranium cartilages at the point of connection with the trabeculae) (Fig. 1D-I). Only a single ethanol-treated wild type sibling (from a large experimental group of *bmp2b* larvae) displayed a misshapen trabecula (Table 1), confirming that our exposure paradigm does not impact wild type larvae as previously described (Bilotta et al., 2004; Everson et al., 2022; McCarthy et al., 2013; Swartz et al., 2014; Zhang et al., 2014).

**Figure 1.**
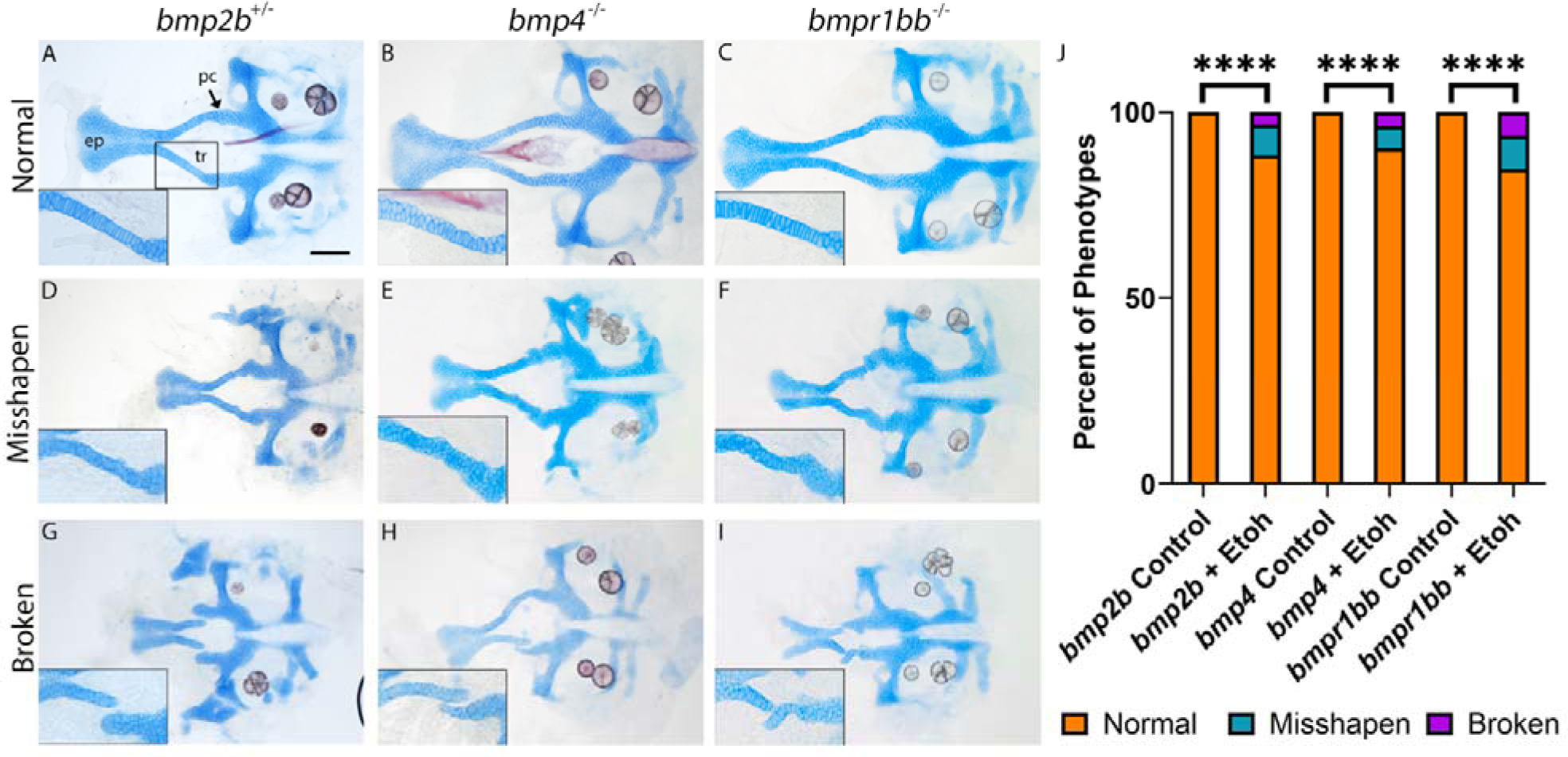
Multiple members of the Bmp pathway exhibit ethanol-induced palate defects. (**A-I**) Flat-mount images of neurocranium of 5 dpf larva. Cartilage is blue and bone is red (anterior to the left, scale bar: 100 μ, ep = ethmoid plate, tr = trabeculae and pc = polar cartilage, large images were captured at 10x, inset is 20x image of the left trabecula). (**J**) total percentage of palate phenotype in ethanol-treated Bmp mutants and controls. (**A-C**) heterozygous *bmp2b* or homozygous *bmp4* or *bmpr1bb* mutant larva develop normal neurocranium. Insets show normal stacking of cells in the trabecula. (**D-I**) Exposure of 1% ethanol results in a range of defects to palate, from misshapen trabecula (**D-F**) to breaks in the trabeculae at the polar cartilages (**G-I**). Percent of phenotypes were quantified and compared for untreated and ethanol-treated larvae for each Bmp component using Fisher’s Exact Test (**** = p <0.0001) (**J**). “n” for each Bmp component listed in Table 1.

**Table 1.**
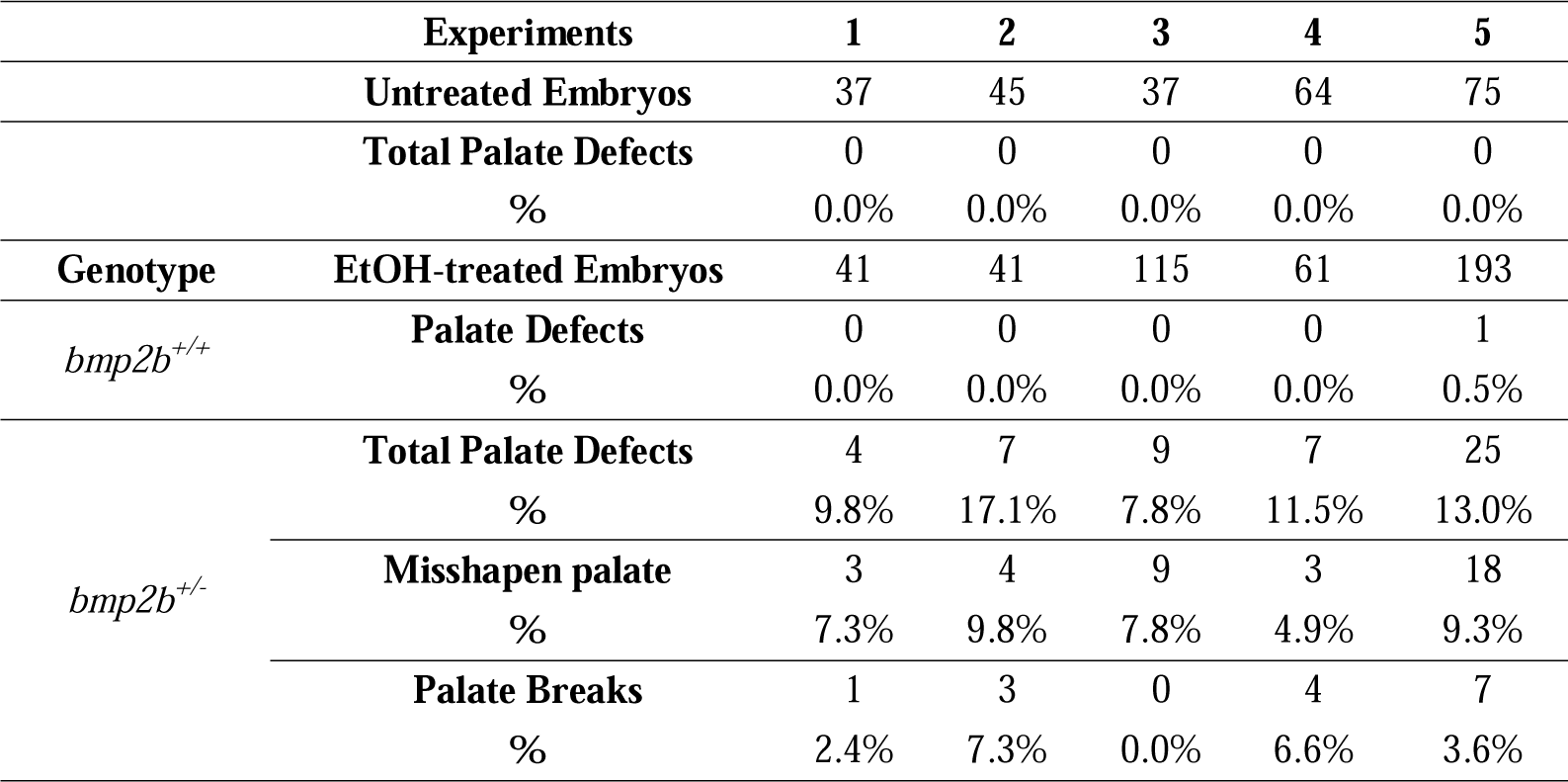
Penetrance of palate defects in ethanol-treated *bmp2b* larvae. Percent of palate defects, both misshapen and broken trabeculae, per genotype, per experiment in ethanol-treated *bmp2b* larvae.

| Experiments |  | 1 | 2 | 3 | 4 | 5 |
| --- | --- | --- | --- | --- | --- | --- |
| Untreated Embryos |  | 37 | 45 | 37 | 64 | 75 |
| Total Palate Defects |  | 0 | 0 | 0 | 0 | 0 |
| % |  | 0.0% | 0.0% | 0.0% | 0.0% | 0.0% |
| Genotype | EtOH-treated Embryos | 41 | 41 | 115 | 61 | 193 |
| *bmp2b*<sup>+/+</sup> | Palate Defects | 0 | 0 | 0 | 0 | 1 |
|  | % | 0.0% | 0.0% | 0.0% | 0.0% | 0.5% |
| *bmp2b*<sup>+/-</sup> | Total Palate Defects | 4 | 7 | 9 | 7 | 25 |
|  | % | 9.8% | 17.1% | 7.8% | 11.5% | 13.0% |
|  | Misshapen palate | 3 | 4 | 9 | 3 | 18 |
|  | % | 7.3% | 9.8% | 7.8% | 4.9% | 9.3% |
|  | Palate Breaks | 1 | 3 | 0 | 4 | 7 |
|  | % | 2.4% | 7.3% | 0.0% | 6.6% | 3.6% |

The expressivity and penetrance of both malformed and broken trabeculae were significant compared to untreated siblings and largely consistent between the different ethanol-treated mutant lines, though the penetrance for both phenotypes was slightly higher in ethanol-treated *bmpr1bb* heterozygous and homozygous mutant larvae (Fig. 1J, Tables 1-3). Interestingly, the penetrance and expressivity for both misshapen and broken trabeculae were largely distributed equally between heterozygous and homozygous mutant larvae for both *bmp4* and *bmpr1bb*, but never observed in wild type siblings (Tables 2 and 3). These results reveal that consistent defects to the palate occur in ethanol-treated zebrafish with Bmp pathway mutations but not in either Bmp mutant or ethanol treatment alone.

**Table 2.** Penetrance of palate defects in ethanol-treated *bmp4* larvae. Percent of palate defects, both misshapen and broken trabeculae, per genotype, per experiment in ethanol-treated *bmp4* larvae.

| Experiments |  | 1 | 2 | 3 | 4 | 5 | 6 |
| --- | --- | --- | --- | --- | --- | --- | --- |
| Untreated Embryos |  | 24 | 41 | 42 | 51 | 44 | 79 |
| Total Palate Defects |  | 0 | 0 | 0 | 0 | 0 | 0 |
| % |  | 0.0% | 0.0% | 0.0% | 0.0% | 0.0% | 0.0% |
| Genotype | EtOH-treated Embryos | 63 | 131 | 38 | 94 | 82 | 174 |
| *bmp4*<sup>+/+</sup> | Palate Defects | 0 | 0 | 0 | 0 | 0 | 0 |
|  | % | 0.0% | 0.0% | 0.0% | 0.0% | 0.0% | 0.0% |
| *bmp4*<sup>+/-</sup> | Total Palate Defects | 4 | 9 | 2 | 5 | 1 | 2 |
|  | % | 6.3% | 6.9% | 5.3% | 5.3% | 1.2% | 1.1% |
|  | Misshapen palate | 1 | 7 | 1 | 1 | 1 | 2 |
|  | % | 1.6% | 5.3% | 2.6% | 1.1% | 1.2% | 1.1% |
|  | Palate Breaks | 3 | 2 | 1 | 4 | 0 | 0 |
|  | % | 4.8% | 1.5% | 2.6% | 4.3% | 0.0% | 0.0% |
| *bmp4*<sup>-/-</sup> | Total Palate Defects | 4 | 7 | 3 | 9 | 2 | 9 |
|  | % | 6.3% | 5.3% | 7.9% | 9.6% | 2.4% | 5.2% |
|  | Misshapen palate | 3 | 6 | 3 | 3 | 1 | 6 |
|  | % | 4.8% | 4.6% | 7.9% | 3.2% | 1.2% | 3.4% |
|  | Palate Breaks | 1 | 1 | 0 | 6 | 1 | 3 |
|  | % | 1.6% | 0.8% | 0.0% | 6.4% | 1.2% | 1.7% |

**Table 3.** Penetrance of palate defects in ethanol-treated *bmpr1bb* larvae. Percent of palate defects, both misshapen and broken trabeculae, per genotype, per experiment in ethanol-treated *bmpr1bb* larvae.

|  | Experiments | 1 | 2 | 3 |
| --- | --- | --- | --- | --- |
|  | Untreated Embryos | 105 | 48 | 154 |
|  | Total Palate Defects | 0 | 0 | 0 |
|  | % | 0.0% | 0.0% | 0.0% |
| Genotype | EtOH-treated Embryos | 82 | 38 | 69 |
| *bmpr1bb*<sup>+/+</sup> | Palate Defects | 0 | 0 | 0 |
|  | % | 0.0% | 0.0% | 0.0% |
| *bmpr1bb*<sup>+/-</sup> | Total Palate Defects | 7 | 0 | 8 |
|  | % | 8.5% | 0.0% | 11.6% |
|  | Misshapen palate | 5 | 0 | 5 |
|  | % | 6.1% | 0.0% | 7.2% |
|  | Palate Breaks | 2 | 0 | 3 |
|  | % | 2.4% | 0.0% | 4.3% |
| *bmpr1bb*<sup>-/-</sup> | Total Palate Defects | 6 | 1 | 7 |
|  | % | 7.3% | 2.6% | 10.1% |
|  | Misshapen palate | 3 | 1 | 3 |
|  | % | 3.7% | 2.6% | 4.3% |
|  | Palate Breaks | 3 | 0 | 4 |
|  | % | 3.7% | 0.0% | 5.8% |

### Upregulation of the Bmp target gene, *GATA3,* does not rescue ethanol-induced palate defects in *bmp4* mutant larvae

Multiple studies have shown that Bmp signals directly to the NCCs and loss of Bmp signaling results in cleft palate (Alexander et al., 2011; Dudas et al., 2004; Ko et al., 2007; Liu, 2005; Swartz et al., 2021, 2011). We and other have shown multiple genes in the NCC are Bmp-dependent showing reduced expression in Bmp knockdown, including *satb2*, *edn1*, *hand2* and *msxe* (Alexander et al., 2011; Sheehan-Rooney et al., 2013). We have previously shown that the NCC transcription factor *gata3* is downstream of Bmp signaling regulating palate formation and that upregulation of human *GATA3* alleviates loss of Bmp signaling during palate development (Swartz et al., 2021). This suggests that upregulating *GATA3* will restore palate formation in ethanol-treated Bmp mutant larvae. From our Bmp mutants, we chose to work on *bmp4* mutant larvae in all subsequent experiments as the mutation of *bmp2b* is weakly dominant (Kishimoto et al., 1997) and *bmpr1bb* is in the WIK background, both of which compound our epistatic analysis with heat shock-induced *gata3*. To evaluate if upregulation of *GATA3* rescues ethanol-induced palate defects, I generated *bmp4^-/-^; hsp:GATA3-GFP* double line. Embryos were treated with ethanol starting at 6 hpf and heat shocked at 24 hpf, as previously performed (Swartz et al., 2021). I observed a low percentage of palate defects in both non-heat shocked and heat shocked controls. I only observed these palate defects in *bmp4* mutants that also carried the *hsp:GATA3-GFP* allele; not in *bmp4* mutant or *hsp:GATA3-GFP* carriers alone. It is possible that the *hsp:GATA3-GFP* allele inserted into a region of the genome that is sensitizing a small percentage of *bmp4* mutants to palate defects in the absence of ethanol, though this is speculation (Fig. 2; Table 4). Ethanol exposure without heat shock resulted in over 20% palate defects, a significant increase over controls. Surprisingly, upregulation of *GATA3* did not rescue ethanol-induced palate defects (Fig. 2; Table 4). In actuality, I observed a significant increase in palate defects in ethanol-treated, heat shocked larvae (Fig 2; Table 4). This suggests that upregulation of *gata3* alone is not sufficient to rescue ethanol-induced palate defects in Bmp mutants.

**Figure 2.**
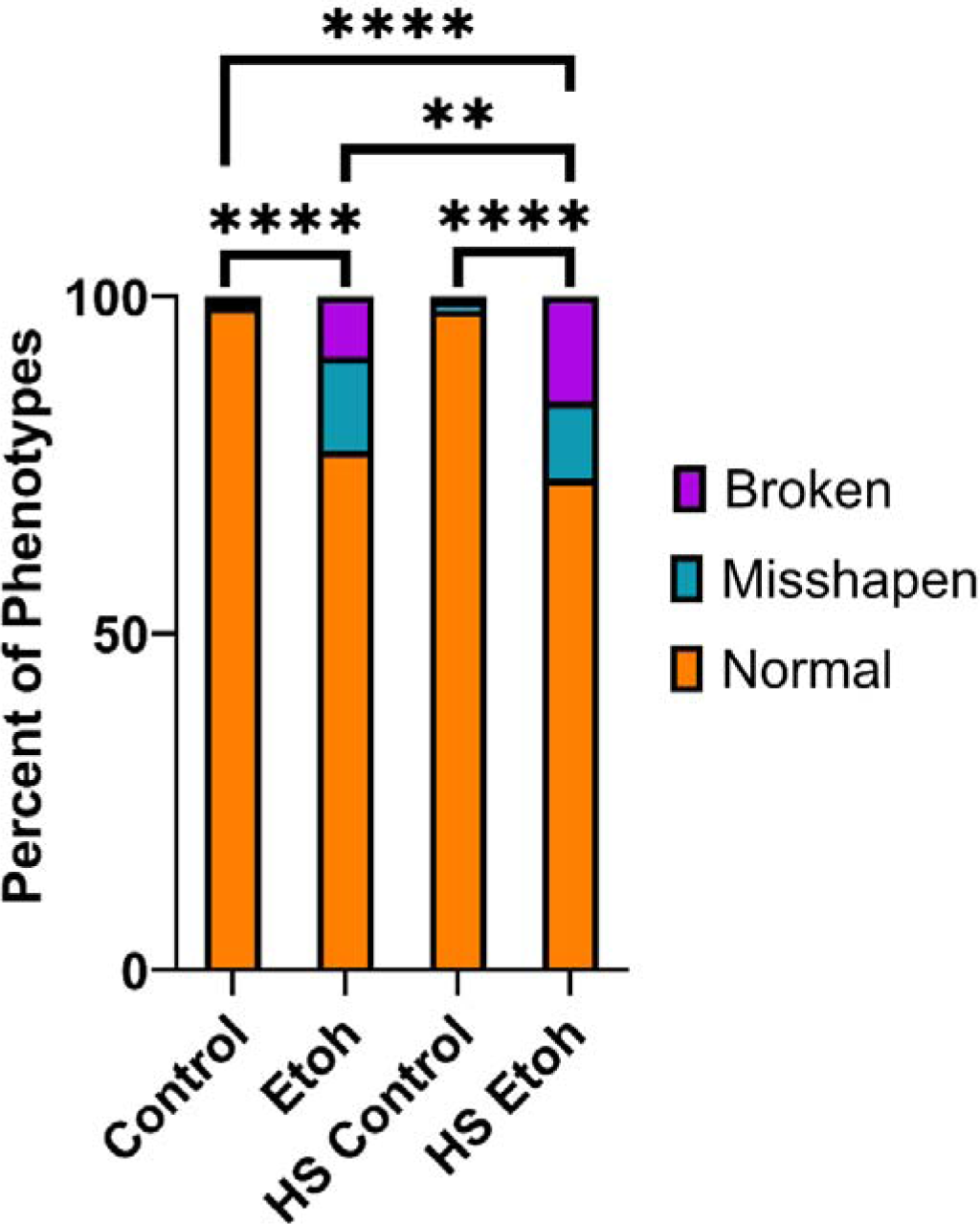
Upregulation of *GATA3* does not rescue ethanol-induced palate defects in *bmp4* mutant larvae. Comparison of palate defects in non-heat shocked control and ethanol-treated *bmp4* mutant larvae to heat shocked control and ethanol-treated *bmp4* mutant larvae. Ethanol exposure results in over 20% of palate defects in *bmp4* mutant larvae. Heat shock induced, upregulation of *GATA3* does not rescue ethanol induced palate defects in *bmp4* mutant larvae. Percent of phenotypes were quantified and compared for untreated and ethanol-treated larvae for each Bmp component using (two-way ANOVA, F = 57.77, p < 0.0001; pairwise comparisons **** = p < 0.0001, ** = p < 0.005). “n” for each treatment group listed in Table 4.

**Table 4.** Percent of palate defects in *hsp:GATA3-GFP* upregulation in *bmp4* mutant larvae. Percent of palate defects, both misshapen and broken trabeculae, per treatment group *bmp4^-/-^; hsp:GATA3-GFP* larvae.

| Experiments | Total Embryos | Palate Defects | Percent | Misshapen palate | Percent | Palate Breaks | Percent |
| --- | --- | --- | --- | --- | --- | --- | --- |
| Control | 198 | 3 | 1.5% | 2 | 1.0% | 1 | 0.5% |
| Ethanol | 201 | 46 | 22.9% | 28 | 13.9% | 18 | 9.0% |
| Heat Shock Control | 287 | 6 | 2.1% | 4 | 1.4% | 2 | 0.7% |
| Heat Shock + Etoh | 218 | 59 | 27.1% | 25 | 11.5% | 34 | 15.6% |

To confirm *gata3* expression is not critical in ethanol-induced palate defects in our Bmp mutants, I asked if *gata3* expression is reduced or lost in ethanol-treated *bmp4* mutant embryos. Our previous work showed that disruption to Bmp signaling either by mutation of the downstream transcription factor, s*mad5*, or genetic knock down via a heat shock inducible dominant negative Bmp receptor results in loss of *gata3* expression in the maxillary domain of the 1^st^ NCC pharyngeal arch (Swartz et al., 2021). To confirm this result, I performed Hybridization Chain Reaction-based (HCR) *in situ* hybridization analysis of *gata3* expression in control ethanol-treated wild type and *bmp4* mutant embryos. My results show that *gata3* expression was not disrupted in both untreated and ethanol-treated *bmp4* mutants compared to controls suggesting that ethanol does not attenuate the Bmp-*gata3* signaling pathway in the NCCs but possibly other aspects of Bmp signaling.

### *bmp4* mutant larvae are sensitive to ethanol from 10-18 hpf leading to palate defects

We have previously shown that Bmp signaling is required in the endoderm from 10-18 hpf for formation of the viscerocranium (Lovely et al., 2016). We have recently shown that our Bmp mutant lines are also ethanol sensitive from 10-18 hpf with exposure to ethanol after 18 hpf failing to lead to any observable developmental defects (Klem et al., 2023 (in revision)). To test the onset of ethanol-induced palate defects, I started our 1% ethanol exposure paradigm on wild type and *bmp4^-/-^* embryos at 10 hpf, 14 hpf, 18 hpf, 22 hpf or 24 hpf and leaving it on out to 5 dpf. My analysis showed that as the onset of ethanol exposure at 10 hpf and 14 hpf lead to consistent palate defects as previously observed (Fig. 4A; Table 5). However, the onset of ethanol exposure at or after 18 hpf did not lead to broad palate defects with only 1% of embryos showing any palate defects compared to 10.3% and 8.8% palate defects at 10 hpf and 14 hpf, respectively (Fig. 4A; Table 5). We have previously shown that ethanol tissue concentrations decrease after 24 hpf compared to exposures starting at 6 hpf (Lovely et al., 2014) suggesting that a higher dose at 24 hpf may elicit palate defects in *bmp4* mutant larvae. To test this, I repeated starting our exposure paradigm at 24 hpf but increased our ethanol exposure concentration from 1% to 1.3%. Again, I did not observe palate defects to *bmp4* mutant larvae when exposure starts at 24 hpf, regardless of dose (Fig 4B; Table 6). Combined, these results suggest that our Bmp mutants are ethanol sensitive from 10-18 hpf, leading to palate defects.

**Figure 3.**
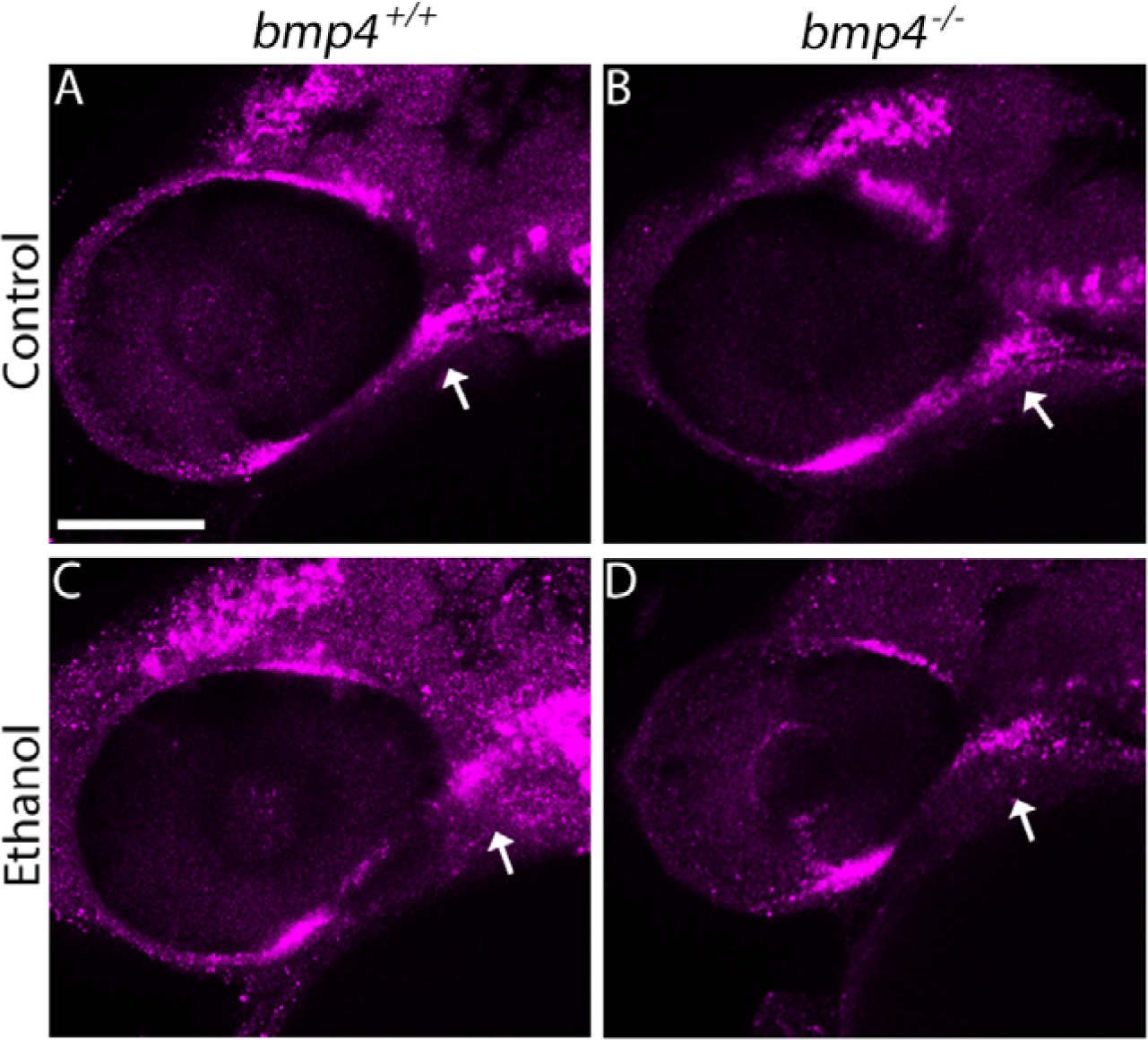
Expression of *gata3* is unaltered in ethanol-treated *bmp4* mutant embryos. (**A-D**) Whole-mount, confocal images of *bmp4* embryos fluorescently labeling *gata3* gene expression at 36 hpf (lateral views, anterior to the left, scale bar: 100 μ). Arrows show normal expression of *gata3* in the maxillary domain of the NCC in untreated and ethanol-treated wild type and *bmp4* mutant embryos as well as ethanol-treated wild type embryos (n = 10 embryos per group).

**Figure 4.**
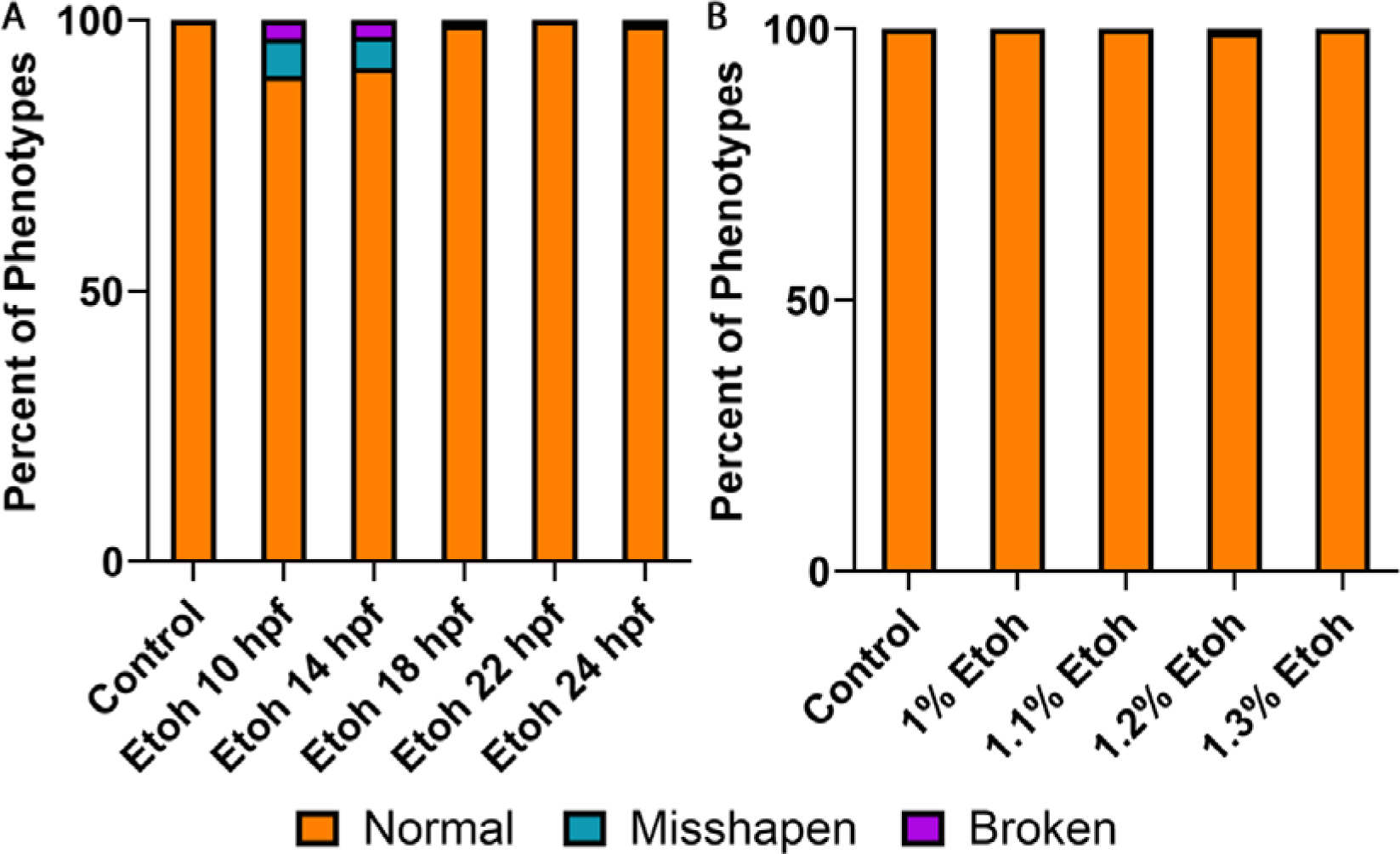
Timing and penetrance of ethanol sensitivity in *bmp4* mutant larvae shows that palate defects occur when exposure is from 10-18 hpf. (**A**) Comparison of palate defects in *bmp4* mutant larvae when ethanol exposure is initiated at 10 hpf, 14 hpf, 18 hpf, 22 hpf or 24 hpf. Ethanol exposure results in approximately 10% of palate defects in *bmp4* mutant larvae when ethanol exposure is initiated at 10 or 14 hpf. Ethanol exposures initiated at 18 hpf or after does not result in palate defects. n” for each treatment group listed in Table 5. (**B**) Comparison of palate defects in *bmp4* mutant larvae with increasing ethanol exposure doses initiated at 24 hpf. Increasing ethanol concentrations, from 1% to 1.3%, initiated at 24 hpf do not result in palate defects. “n” for each treatment group listed in Table 6.

**Table 5.** Timing of ethanol-induced palate defects in *bmp4* mutant larvae. Percent of palate defects, both misshapen and broken trabeculae, per ethanol exposure initiation time point in *bmp4* larvae.

| Experiments | Untreated Embryos | EtOH-treated Embryos | Palate Defects | Percent | Misshapen palate | Percent | Palate Breaks | Percent |
| --- | --- | --- | --- | --- | --- | --- | --- | --- |
| Control | 97 |  | 0 | 0.0% | 0 | 0.0% | 0 | 0.0% |
| E10 |  | 174 | 18 | 10.3% | 12 | 6.9% | 6 | 3.4% |
| E14 |  | 193 | 17 | 8.8% | 11 | 5.7% | 6 | 3.1% |
| E18 |  | 192 | 2 | 1.0% | 1 | 0.5% | 1 | 0.5% |
| E22 |  | 197 | 0 | 0.0% | 0 | 0.0% | 0 | 0.0% |
| E24 |  | 194 | 2 | 1.0% | 1 | 0.5% | 1 | 0.5% |

**Table 6.** Ethanol dose response of palate defects in *bmp4* mutant larvae after 24 hpf. Percent of palate defects, both misshapen and broken trabeculae, per ethanol exposure concentration in *bmp4* mutant larvae.

| Experiments | Untreated Embryos | EtOH-treated Embryos | Palate Defects | Percent | Misshapen palate | Percent | Palate Breaks | Percent |
| --- | --- | --- | --- | --- | --- | --- | --- | --- |
| Control | 138 |  | 0 | 0.0% | 0 | 0.0% | 0 | 0.0% |
| E 1% |  | 165 | 0 | 0.0% | 0 | 0.0% | 0 | 0.0% |
| E 1.1% |  | 159 | 0 | 0.0% | 0 | 0.0% | 0 | 0.0% |
| E 1.2% |  | 165 | 1 | 0.6% | 1 | 0.6% | 0 | 0.0% |
| E 1.3% |  | 159 | 0 | 0.0% | 0 | 0.0% | 0 | 0.0% |

### Blocking Bmp with the small chemical inhibitor dorsomorphin from 10-18 hpf does not disrupt *gata3* expression

To confirm that 10-18 hpf is the ethanol-sensitive time window for palate defects, I examined multiple time windows of Bmp knockdown with DM. We have previously shown that zebrafish embryos treated with DM from 10-18 hpf lose Bmp signaling responses in the pharyngeal endoderm leading to jaw defects (Lovely et al., 2016). Here, I confirm that DM treatment starting at 10 hpf and lasting out to 18 hpf leads to palate defects with the shortest window of exposure starting at 10 hpf leading to palate defects being 2 hours, 10-12 hpf (Fig. 5A; Table 7). To identify the latest starting point in which Bmp signaling could be blocked and still lead to palate defects, I initiated our exposure windows every hour starting at 14 hpf and ending at 22 hpf, leaving DM on until 30 hpf. I chose to start at 14 hpf as that is the last window in which initiation of ethanol exposure still results in palate defects. DM exposure from 14-30 hpf leads to 100% penetrance of palate defects with the penetrance slowly decreasing at the initiation time points of 15 hpf, 16 hpf and 17 hpf (Fig. 5B; Table 7). The percent of palate defects sharply declines starting at 18 hpf with initiation of DM exposure at 20 hpf and later failing to generate any palate defects (Fig. 5B; Table 7). This suggests that blocking Bmp signaling from 10-18 hpf is the critical time window for palate development. To further confirm this, I examined *gata3* expression in embryos treated with DM from 10-18 hpf. I did not see any change in *gata3* expression when Bmp signaling is blocked (Fig. 5C&D). This mirrors what I observed in our ethanol exposure windows analysis and suggests that Bmp signaling is required from 10-18 hpf for palate development and that either ethanol treatment of Bmp mutants or blocking Bmp signaling with DM disrupts palate development independent of *gata3*.

**Figure 5.**
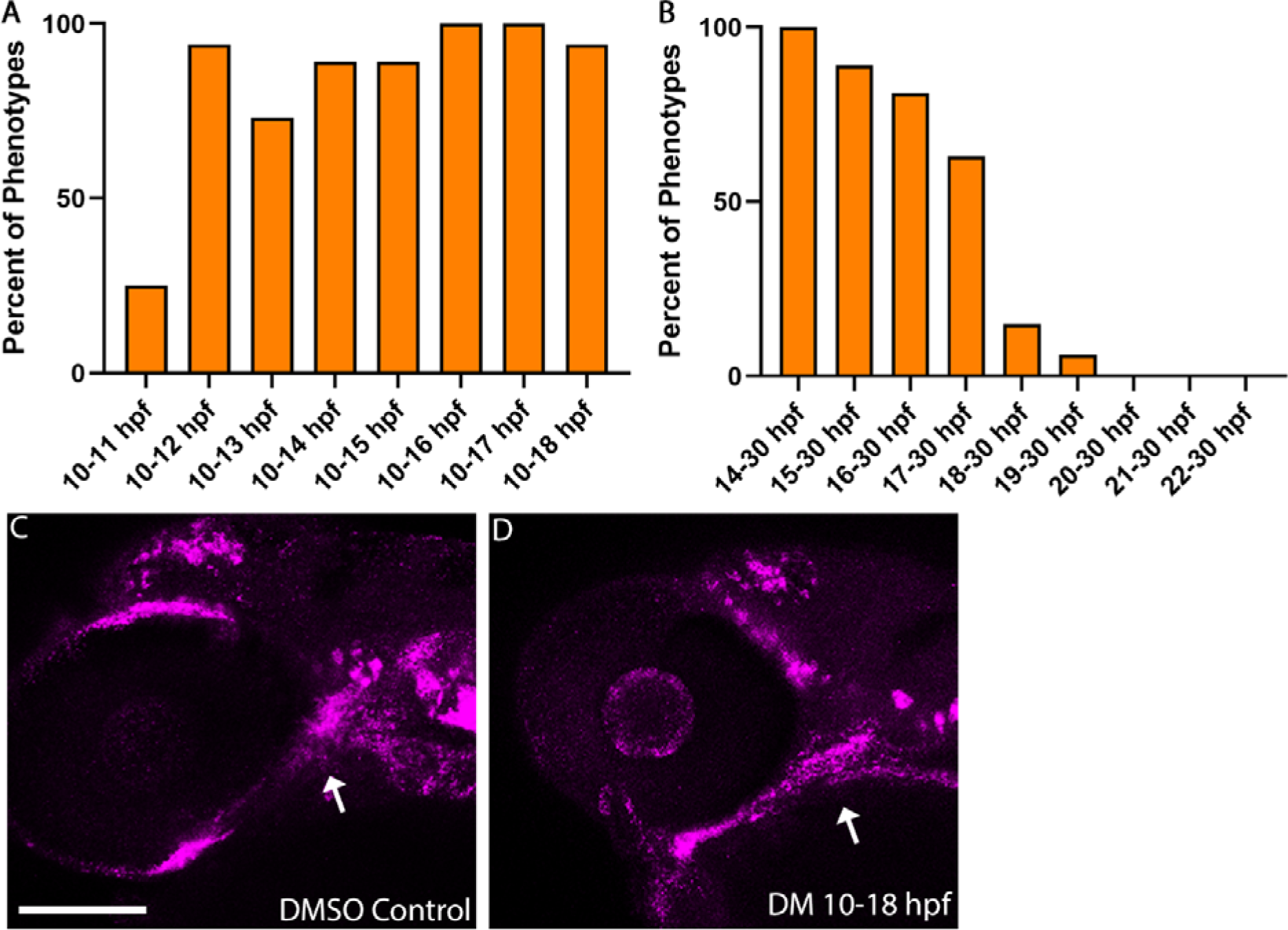
Dorsomorphin-treated larvae show palate defects but normal *gata3* expression when treated between 10 and 18 hpf. (**A**) Comparison of palate defects in DM-treated larvae with expanding DM exposure window from 10-11 hpf out to 10-18 hpf. We observed a large number of palate defects with a DM exposure window as short as 2 hours, 10-12 hpf with increasing duration of exposure resulting in consistent palate defects. (**B**) Comparison of palate defects in DM-treated larvae with increasingly later initiation time points of DM, 14 hpf (consistent with the latest exposure initiation of ethanol at a higher penetrance (Fig. 4A) out to 30 hpf. Consistent with the ethanol sensitivity exposure window, the percentage of palate defects decreases drastically when DM exposure is initiated at 18 hpf. “n” for each treatment group listed in Table 7. (**C&D**) Whole-mount, confocal images of DM-treated (10-18 hpf) or DMSO control embryos fluorescently labeling *gata3* gene expression at 36 hpf (lateral views, anterior to the left, scale bar: 100 μm). Arrows show normal expression of *gata3* in the maxillary domain of the NCC in DMSO and DM-treated embryos as well as ethanol-treated wild type embryos (n = 7 embryos per treatment group).

**Table 7.** Timing of DM induced palate defects. Percent of palate defects per DM exposure window duration and initiation in WT larvae.

| **Exposure Window** | **Untreated Embryos** | **DM-treated Embryos** | **Palate Defects** | **Percent** |
| --- | --- | --- | --- | --- |
| **Control** | 20 |  | 0 | 0.0% |
| **10-11 hpf** |  | 16 | 4 | 25.0% |
| **10-12 hpf** |  | 17 | 16 | 94.1% |
| **10-13 hpf** |  | 15 | 11 | 73.3% |
| **10-14 hpf** |  | 19 | 17 | 89.5% |
| **10-15 hpf** |  | 18 | 16 | 88.9% |
| **10-16 hpf** |  | 19 | 19 | 100.0% |
| **10-17 hpf** |  | 14 | 14 | 100.0% |
| **10-18 hpf** |  | 17 | 16 | 94.1% |
| **Control** | 19 |  | 0 | 0.0% |
| **14-30 hpf** |  | 17 | 17 | 100% |
| **15-30 hpf** |  | 19 | 17 | 89% |
| **16-30 hpf** |  | 15 | 13 | 81% |
| **17-30 hpf** |  | 16 | 10 | 63% |
| **18-30 hpf** |  | 17 | 2 | 15% |
| **19-30 hpf** |  | 17 | 1 | 6% |
| **20-30 hpf** |  | 18 | 0 | 0% |
| **21-30 hpf** |  | 17 | 0 | 0% |
| **22-30 hpf** |  | 17 | 0 | 0% |

## Discussion

Palate formation is driven by a distinct set of cell behaviors in the NCCs with disruption of these behaviors leading to defects formation of the palate (Greene and Pisano, 2010; Mork and Crump, 2015; Siismets and Hatch, 2020). Proper genetic regulation of these behaviors is critical for palate formation (Cobourne, 2004; Şahin Uysal et al., 2023; Stanier, 2004). Numerous genes and signaling pathways, which when attenuated either genetically and/or environmentally, drive the cell behaviors underlying palate formation (Bemquerer et al., 2022; Dixon et al., 2011; Marazita, 2023; Narumi et al., 2020). Prenatal ethanol exposure, defined as Fetal Alcohol Spectrum Disorders (FASD) describe a highly variable set of phenotypes to the facial skeleton, including the palate (Eberhart and Parnell, 2016; Everson and Eberhart, 2023). Ethanol-sensitive genetic loci are major drivers of FASD and provide insight into the signaling events potentially disrupted in FASD (Eberhart and Parnell, 2016; Lovely, 2020; McCarthy and Eberhart, 2014). Here we show that mutation in multiple components of the Bmp signaling pathway sensitize zebrafish embryos to ethanol-induced palate defects with mutations in *bmp2b*, *bmp4* and *bmpr1bb*, leading to malformations of the trabeculae. These defects divide into two main phenotypes, misshapen trabeculae where the shape of the cells is altered, failing to stack out and form the smooth bilateral rod structures. The second is breaks in the trabeculae at the polar cartilages of the posterior neurocranium. Both the penetrance and expressivity of these defects were consistent across all three Bmp mutant lines with both the misshapen and broken trabeculae phenotypes were observed in heterozygous homozygous mutant carriers but never wild type siblings. This strongly suggests a consistent mechanism driving these defects in Bmp mutants. However, future work will be needed to examine the detailed cellular phenotypes in these ethanol-treated Bmp mutants.

Multiple studies have demonstrated a major role for Bmp in palate formation with loss of Bmp signaling leading to cleft palate (Alexander et al., 2011; Dudas et al., 2004; Ko et al., 2007; Liu, 2005; Swartz et al., 2021, 2011). We have previously shown that NCC specific, Bmp signaling is necessary for palate morphogenesis in a *gata3* dependent manner (Swartz et al., 2021, 2011). Mutations in human *GATA3,* which result in the highly variable hypoparathyroidism, sensorineural deafness and renal dysplasia (HDR) syndrome, is also associated with palatal defects (Bilous et al., 1992; Van Esch et al., 2000). Zebrafish *gata3* mutants lose a specific region of the palatal skeleton, the trabeculae, similar to loss of Bmp signaling (Swartz et al., 2021, 2011). Upregulation of human *GATA3* in zebrafish rescues these Bmp dependent palate defects (Swartz et al., 2021). Surprisingly, overexpression of *GATA3* in our ethanol-treated Bmp mutants does not rescue ethanol induced palate defects and *gata3* expression was unaltered in our ethanol treated *bmp4* mutants. This contradicts our previous studies of the role of Bmp-*gata3* in palate formation. However, we have previously shown that gene-ethanol interactions can exhibit similar phenotypes as untreated mutants, but through novel mechanisms that may not be readily predicted from the mutation alone (McCarthy and Eberhart, 2014; Sidik et al., 2021; Swartz et al., 2020, 2014). Ultimately, this suggest that the Bmp-ethanol interaction disrupts palate development through a unique mechanism, independent of *gata3*.

We have previously shown that Bmp signaling is required in the pharyngeal endoderm for proper endoderm morphogenesis and subsequent facial development form 10-18 hpf (Lovely et al., 2016). We have recently shown that Bmp-dependent jaw development is also ethanol sensitive from 10-18 hpf suggesting this narrower time window drives ethanol-induced palate defects as well (Klem et al., 2023 (in revision)). I assessed multiple time windows and saw that the exposure paradigm of 10-18 hpf that drove jaw defects also drives palate defects. Any exposure at or after 18 hpf did not result in palate defects, even at higher doses of ethanol, similar to what we observed for ethanol-induced jaw defects (Klem et al., 2023 (in revision)). In our previous study, we went on to show that Bmp-ethanol interactions when exposed from 10-18 hpf disrupt anterior endoderm morphogenesis, increasing the size of the anterior endoderm, leading to jaw defects (Klem et al., 2023 (in revision)). Previous work has shown that the anterior endoderm is necessary to induce signaling factor expression in the oral ectoderm, in particular *fgf8a* (Balczerski et al., 2012; Haworth et al., 2007, 2004). We show that ethanol-induced increase anterior endoderm size leads to expanded expression of *fgf8a* (Klem et al., 2023 (in revision)). This is consistent with previous work showing that malformations of the anterior endoderm in the sphingosine signaling mutant *s1pr2* disrupt oral ectoderm expression domains leading to jaw defects (Balczerski et al., 2012). Interestingly, the *s1pr2* mutants also have palate defects (Balczerski et al., 2012), supporting that defects to anterior endoderm-oral ectoderm signaling disrupt both palate and jaw development. Overall, these observations indicate that our Bmp-ethanol interactions disrupt an anterior endoderm-oral ectoderm-signaling axis that is necessary for formation of the anterior neurocranium.

While we have shown that Bmp-ethanol interactions disrupt epithelial development leading to jaw and palate defects, the mode of action of ethanol is still unknown. We recently quantified Corrected Total Fluorescence of the Bmp response in the pharyngeal arches using the Bmp responsive transgenic line, *BRE:mKO2* that labels active, canonical p-Smad-based, Bmp signaling (Collery and Link, 2011; Klem et al., 2023 (in revision)). We employed this approach as it gave us the ability to quantify tissue specific impacts of ethanol on Bmp signaling. From this, we observed loss of endodermal-specific Bmp signaling responses in our untreated Bmp mutants while Bmp signaling in other tissues remained intact. Ethanol, however, does not disrupt Bmp signaling in these other tissues in either Bmp mutants or their wild type siblings, suggesting that ethanol is acting independent of canonical Bmp signaling. Several possibilities exist explaining these results: 1) ethanol disrupts non-canonical Bmp signaling events, i.e. LIMK/cofilin mediated non-canonical BMP receptor signaling (Park and Gumbiner, 2012). 2) ethanol disrupts function of one or several downstream Bmp target(s). We observed similar results in our previous analyses of *pdgfra*-ethanol interactions where ethanol acted at the level of mTOR, downstream of *pdgfra* (McCarthy et al., 2013). 3) ethanol disrupts signaling events / cellular processes in the endoderm that are independent yet converge with the role of Bmp signaling in the endoderm. 4) ethanol may act on tissues other than the endoderm, disrupting cellular events in other tissues (i.e. the NCCs). Ultimately, while ethanol does not directly disrupt canonical Bmp signaling, ethanol does potentiate Bmp mutation to elicit defects in jaw and palate formation. Future analyses will be needed to determine ethanol-sensitive mechanisms that sensitize Bmp mutants to these craniofacial defects.

Ethanol is incredibly pleiotropic, impacting numerous cell processes (Eberhart and Parnell, 2016; Lovely, 2020). Ethanol has been shown to increase NCC specific cell death in multiple model systems (Smith et al., 2014). We have previously shown that loss of *plk1* sensitizes embryos to ethanol-induced cell death with loss of neural tissues and the entire facial skeleton, while loss of *pdgfra* leads to increased ethanol-induced cell death in the NCCs that give rise to the zebrafish palate (McCarthy et al., 2013). It is possible that Bmp mutation potentiates the embryos to ethanol-induced NCCs death leading to the facial defects we observe in both the palate and the jaw. Our future analyses will examine ethanol-induced cell death in the in NCC as well as the endoderm and surrounding epithelial tissues. It is also possible that ethanol may impact epithelial development directly converging the role of Bmp signaling in endoderm morphogenesis and tissue interactions. Any change in these endoderm-oral ectoderm-NCC interactions could lead to the palate defects we observe in our ethanol-treated Bmp mutants. While our data supports this hypothesis, more work needs to be done to examine the impact of ethanol on the cell behaviors and signaling interactions of the endoderm-oral ectoderm-NCC signaling axis and how these events disrupt subsequent facial development.

Combined, my results show that Bmp-ethanol interactions disrupt palate formation in manner not predicted by loss of Bmp signaling alone. While Bmp signaling is required in the NCCs for *gata3* expression and subsequent palate development, I show that ethanol impacts earlier events in Bmp mutants, independent of NCC specific *gata3*. This demonstrates that despite our increased understanding of the genetics of development, much remains to be learned of the impact of ethanol on the signaling mechanisms underlying FASD (Lovely, 2020). Here, I expand our current understanding of Bmp-ethanol interactions in FASD, and, ultimately, provide a conceptual framework for future FASD studies of complex signaling pathways driving facial development.

## Acknowledgements

The author would like to thank Kevin Kump for zebrafish animal care and husbandry. I thank Dr. Jim Amatruda for the providing the *bmpr1bb* zebrafish line. I also thank Dr. Duygu Özpolat and Dr. Ryan Hull for providing script for HCR probe design.

## Author Contributions

CBL conceived the project, designed all studies, conducted all experiments, analyzed all data, generated all figures, wrote the manuscript.

